# Highly Efficient, Massively-Parallel Single-Cell RNA-Seq Reveals Cellular States and Molecular Features of Human Skin Pathology

**DOI:** 10.1101/689273

**Authors:** Travis K Hughes, Marc H Wadsworth, Todd M Gierahn, Tran Do, David Weiss, Priscilla R. Andrade, Feiyang Ma, Bruno J. de Andrade Silva, Shuai Shao, Lam C Tsoi, Jose Ordovas-Montanes, Johann E Gudjonsson, Robert L Modlin, J Christopher Love, Alex K Shalek

## Abstract

The development of high-throughput single-cell RNA-sequencing (scRNA-Seq) methodologies has empowered the characterization of complex biological samples by dramatically increasing the number of constituent cells that can be examined concurrently. Nevertheless, these approaches typically recover substantially less information per-cell as compared to lower-throughput microtiter plate-based strategies. To uncover critical phenotypic differences among cells and effectively link scRNA-Seq observations to legacy datasets, reliable detection of phenotype-defining transcripts – such as transcription factors, affinity receptors, and signaling molecules – by these methods is essential. Here, we describe a substantially improved massively-parallel scRNA-Seq protocol we term Seq-Well S^3 (“Second-Strand Synthesis”) that increases the efficiency of transcript capture and gene detection by up to 10- and 5-fold, respectively, relative to previous iterations, surpassing best-in-class commercial analogs. We first characterized the performance of Seq-Well S^3 in cell lines and PBMCs, and then examined five different inflammatory skin diseases, illustrative of distinct types of inflammation, to explore the breadth of potential immune and parenchymal cell states. Our work presents an essential methodological advance as well as a valuable resource for studying the cellular and molecular features that inform human skin inflammation.

## INTRODUCTION

Although a nascent technology, single-cell RNA-sequencing (scRNA-Seq) has already helped define, at unprecedented resolution, the cellular composition of many healthy and diseased tissues (Klein et al., 2015; Macosko et al., 2015; Montoro et al., 2018; Ordovas-Montanes et al., 2018; Vento-Tormo et al., 2018). The development of high-throughput methodologies has been crucial to this process, empowering the characterization of increasingly complex cellular samples. Unfortunately, current scRNA-Seq platforms typically demonstrate an inverse relationship between the number of cells that can be profiled at once and the amount of biological information that can be recovered from each cell. As a result, one must choose between quantity and quality – and thus comprehensiveness and fidelity – or alternatively employ two distinct approaches in parallel (Tabula Muris Consortium et al., 2018). Indeed, inefficiencies in transcript capture among massively-parallel methods have limited our ability to resolve the distinct cell states that comprise broad cell types (Braga et al., 2019), as well as their essential molecular attributes and often lowly-expressed molecular features, such as transcription factors, affinity receptors, and signaling molecules (**Figure 1A**).

**Figure 1.**
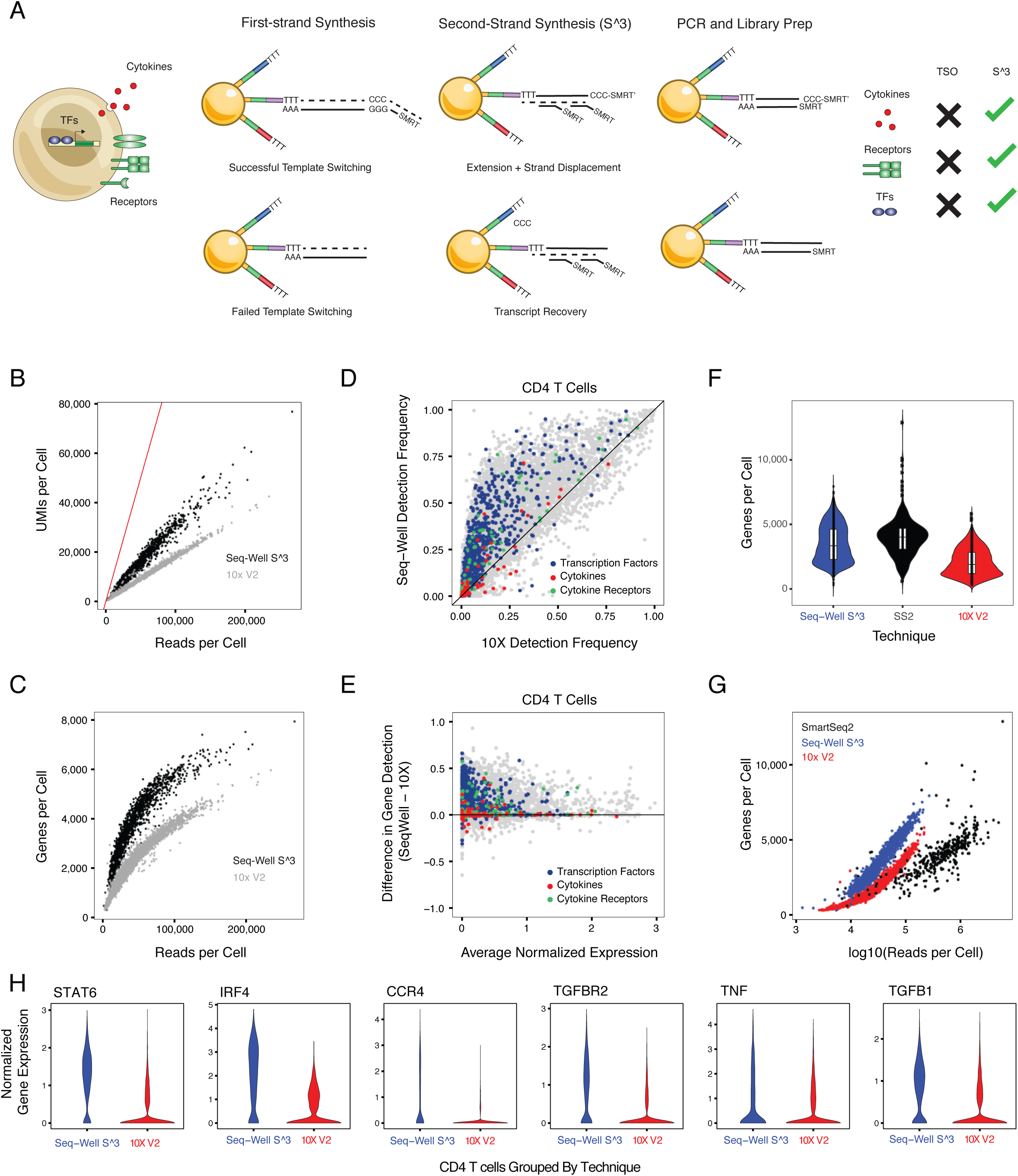
Overview of Second Strand Synthesis (S^3) **A.** Conceptual illustration of the molecular features that define immune phenotypes – including transcription factors, cytokines and receptors – as well as the Seq-Well second-strand synthesis method (Seq-Well S^3) and how it improves detection of key genes and transcripts. **B.** Scatterplot showing differences in per-cell transcript capture (y-axis) as a function of aligned reads per cell (x-axis) between 10x Genomics v2 (10x v2, grey) and Seq-Well S^3 (black). Red line indicates uniform line where transcripts per cell and aligned reads would be equivalent. **C.** Scatterplot shows the differences in per-cell gene detection (y-axis) as a function of aligned reads per cell (x-axis) between 10x v2 (grey) and Seq-Well S^3 (black). **D.** Scatterplot comparing gene detection rates in CD4^+^ T cells between 10x v2 (x-axis) and Seq-Well S^3 (y-axis). Black line indicates point of equivalence in gene detection frequency between methods. Colors correspond to classes of genes including transcription factors (blue), cytokines (magenta), and receptors (green). **E.** Scatterplot comparing gene detection frequency (y-axis) between Seq-Well S^3 (positive values) and 10x v2 (negative values) as a function of the aggregate expression levels (log(scaled UMI + 1)) of an individual gene (x-axis). Black line indicates point of equivalence in gene detection frequency between methods. Colors correspond to classes of genes including transcription factors (blue), cytokines (magenta), and receptors (green). **F.** Violin plot (boxplots median +-quartiles) showing the distribution of per-cell transcript capture for Seq-well S^3 (blue; n = 1,485), 10x v2 (red; n = 2995), and Smart-Seq2 (black, n = 382). **G.** Scatterplot showing the relationship between aligned reads and genes detected per cell between Seq-Well S^3 (blue), 10x v2 (red) and Smart-Seq2 (black) in sorted PBMC CD4^+^ T cells. **H.** Violin plots showing the distribution of normalized expression values for select transcription factors, cytokines and cytokine receptors between Seq-Well S^3 and 10x v2.

Improving the fidelity of these methodologies is particularly important for resolving differences within heterogeneous populations of immune cells like lymphocytes and myeloid cells (Villani et al., 2017). Here, subtle differences in surface receptor, transcription factor and/or cytokine expression can profoundly impact cellular function, particularly in the setting of human pathology (Puel et al., 1998). Enhancing data quality in high-throughput scRNA-Seq would facilitate a greater appreciation of the underlying molecular features that describe such cellular variation. Similarly, it would ease integration with legacy datasets that often rely on lowly-expressed biomarkers, such as transcription factors, that are false-negative prone to discriminate subsets of cells.

Most high-throughput scRNA-Seq methods currently rely on early barcoding of cellular contents to achieve scale. Typically, these techniques recover single-cell transcriptomes for thousands of cells at once by leveraging reverse-emulsion droplets or microwells to isolate individual cells with uniquely barcoded poly-dT oligonucleotides which can then capture and tag cellular mRNAs during reverse transcription (Prakadan et al., 2017). Afterward, an additional priming site is added to the 3’ end of the synthesized cDNA to enable PCR-based amplification of all transcripts using a single primer (whole transcriptome amplification, WTA). A number of techniques have been described to add this second priming site (Sasagawa et al., 2013; Shishkin et al., 2015). The most common uses the terminal transferase activity of certain reverse transcription enzymes to facilitate a “template-switch” from the original mRNA to a second defined oligonucleotide handle (Picelli et al., 2013). While simple to implement, this process has the potential to be highly inefficient, leading to the loss of molecules that have been captured and converted to cDNA but not successfully tagged with a secondary PCR priming site (**Figure 1A** and **S1A**) (Islam et al., 2012; Kapteyn et al., 2010; Zajac et al., 2013).

To overcome these limitations, we have developed a new massively-parallel scRNA-Seq protocol we call Seq-Well S^3 (for “Second-Strand Synthesis”). Seq-Well S^3 increases the efficiency of the second PCR handle addition by amending it through a randomly-primed second-strand synthesis after reverse transcription (**Figure 1A**). Working with cell lines and peripheral blood mononuclear cells (PBMCs), we demonstrate that Seq-Well S^3 enables significant improvements in transcript and gene capture across sample types, facilitating studies of complex immune tissues at enhanced resolution (**Figures 1, S1**, and **S2**).

To illustrate the utility of S^3, we apply it to generate a resource of single-cell transcriptional states spanning multiple inflammatory skin conditions. Skin represents the largest barrier tissue in the human body and is comprised of numerous specialized cell-types that help maintain both immunological and physical boundaries between our inner and outer worlds (Kabashima et al., 2019). The dermis and epidermis – the two primary compartments of human skin – play complementary roles in tissue structure and function (**Figure 2A**) (Kabashima et al., 2019). The epidermis consists primarily of keratinized epithelial cells, which provide a physical barrier to the outside world; the dermis, meanwhile, provides structural support for the skin, with fibroblasts producing collagen and elastin fibrils along with the other components of the extracellular matrix. Crucially, within the cellular ecosystem of human skin, there are numerous tissue-resident immune and parenchymal cells essential to homeostatic barrier function. Using Seq-Well S^3, we examine the cellular composition of normal skin and altered cellular phenotypes in multiple inflammatory skin conditions, including acne, alopecia areata, granuloma annulare, leprosy and psoriasis. With conditions that span autoimmune (alopecia), autoinflammatory (psoriasis), reactive (acne), and granulomatous (granuloma annulare and leprosy) inflammation, we uncover a diverse spectrum of immune and parenchymal cellular phenotypes, as well as their molecular features, across multiple inflammatory skin conditions. Overall, our work presents an essential methodological advance as well as a critical resource for understanding how diverse inflammatory responses can impact a single tissue and the range of cellular phenotypes that are possible upon perturbation.

**Figure 2.**
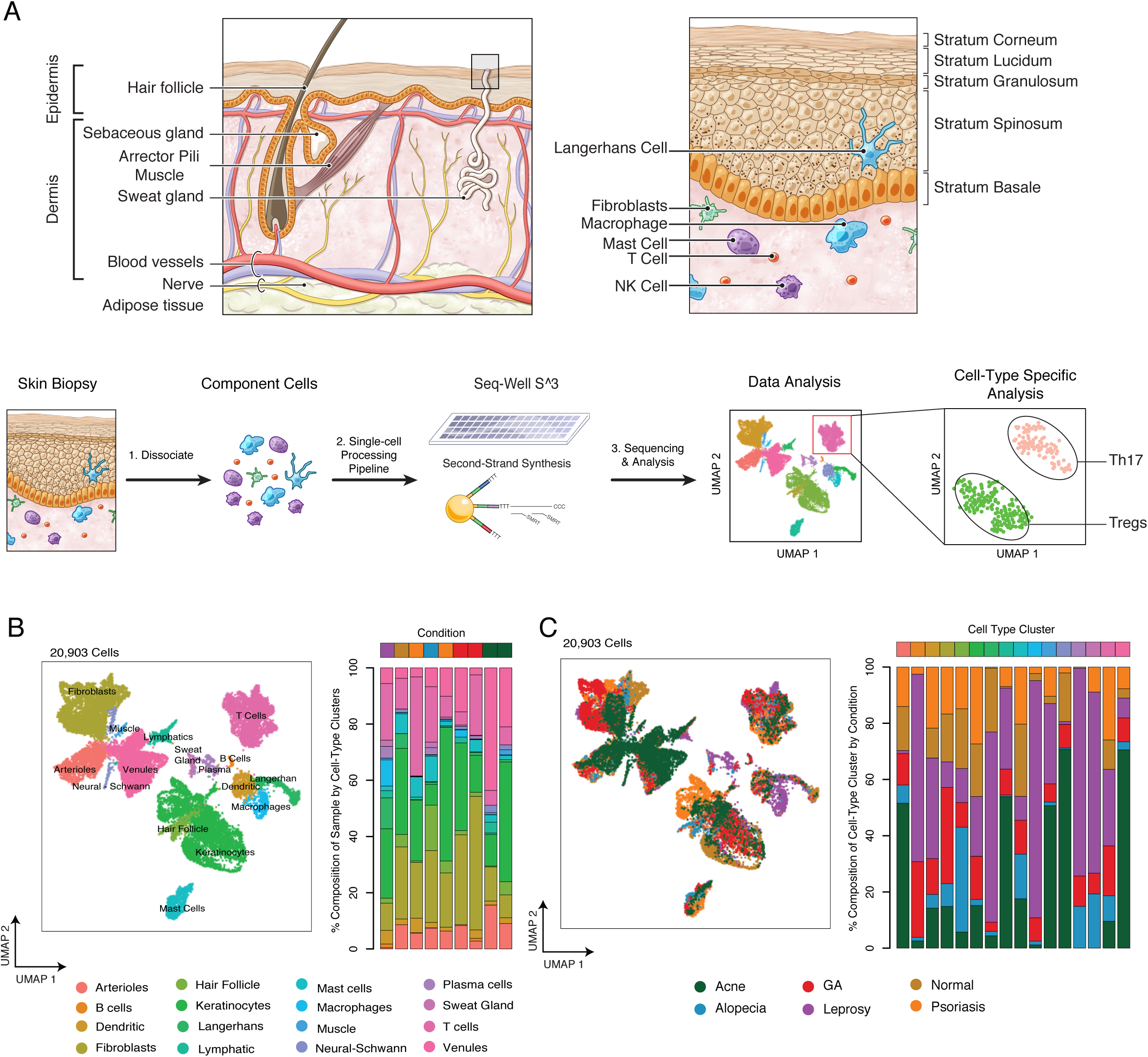
Cell Types Recovered across Inflammatory Skin Conditions. **A.** (Top-Left) Diagram illustrating the anatomic organization and major features of human skin. (Top-Right) Cell-type composition of the epidermis and dermis. (Bottom) Sample processing pipeline used to generate a collection cellular states across skin inflammation. **B.** (Left) UMAP plot for 20,308 cells colored by cell-type cluster. (Right) Stacked barplot showing the cell-type composition for each of the nine skin biopsies. **C.** (Left) UMAP plot for 20,308 cells colored by inflammatory skin condition. (Right) Stacked barplot showing the proportion of cells from each skin condition within phenotypic clusters.

## RESULTS

### Second-Strand Synthesis (S^3) Leads to Improved Transcript Capture and Gene Detection

We hypothesized that use of “template-switching” to append a second PCR handle during reverse transcription might limit the overall recovery of unique transcripts and genes from individual cells in some massively-parallel scRNA-Seq methods such as Seq-Well and Drop-Seq (Gierahn et al., 2017; Macosko et al., 2015). Thus, we incorporated a randomly primed second-strand synthesis following first-strand cDNA construction (**Figures 1A** and **S1A**). Briefly, after reverse transcription, barcoded mRNA capture beads are washed with 0.1 molar sodium hydroxide to remove attached RNA template strands and then a random second-strand synthesis is performed to generate double-stranded cDNA labeled on one end with the SMART sequence and its reverse complement on the other (**Figure 1A** and **S1A**) (Picelli et al., 2013, 2014).

To examine the effectiveness of Seq-Well S^3 and optimize its performance, we first tested a number of conditions using cell lines (**Figure S1B**). In these experiments, we observed that S^3 led to marked improvements in library complexity (Seq-Well V1: 0.22 transcripts/ aligned read, Seq-Well S^3: 0.68 transcripts/ aligned read) and was able to function in the absence of a template switching oligo (TSO); Seq-Well V1, meanwhile, failed to generate appreciable product without a TSO (**Figure S1B-D**). In species-mixing experiments using HEK293 (human) and NIH-3T3 (mouse) cell lines, the use of the S^3 protocol resulted in significant increases in the numbers of unique transcripts captured and genes detected per cell compared to our original protocol for Seq-Well (P < 0.05, Mann-Whitney U Test; **Figure S1C**).

To fully understand how S^3 would perform on more challenging primary cells, we next applied it to human PBMCs (**Figure S1C** and **S2**), benchmarking against our original Seq-Well protocol as well as a commercial technology (10X genomics, V2 chemistry; hereafter 10x v2). For these comparisons, we down-sampled all resulting data to an average of 42,000 reads per cell to account for differences in sequencing depth across technologies. Critically, Seq-Well S^3 resulted in significant improvements in the complexity of our sequencing libraries compared to 10x v2 as determined by the number of transcripts and genes detected at matched read depth (P < 0.05, Mann-Whitney U Test & Linear Regression; **Figure 1B-C**). To confirm that these overall improvements were not driven by changes in the relative frequencies of different cell types captured by each technology, we also examined each subset independently (**Figure S2A-B**). For each cell type detected, we observed significant increases in the numbers of transcripts captured and genes detected using S^3 for each pairwise comparison between techniques (P < 0.05, Mann-Whitney U Test; CD4^+^ T cells, Seq-Well V1: 1,044 ± 62.3 UMIs/cell; 10x v2: 7,671 ± 103.9 UMIs/cell; Seq-Well S^3: 13,390 ± 253.4 UMIs/cell; Mean ± Standard Error of the Median (SEM); **Figure S2**). Both Seq-Well S^3 and 10x v2 displayed increased sensitivity for transcripts and genes relative to Seq-Well v1, but Seq-Well S^3 showed the greatest efficiency (defined as genes recovered at matched read depth) to detect genes for each cell type (**Figure 1D-E**; **Figure S2**).

We sought to further understand whether these improvements resulted in enhanced detection of biologically relevant genes typically under-represented in high-throughput single-cell sequencing libraries (Tabula Muris Consortium et al., 2018). Importantly, genes that were differentially detected (i.e., higher in S^3) within each cell type include numerous transcription factors, cytokines and cell-surface receptors (**Figure 1D-E**). For example, among CD4^+^ T cells, we observe significantly increased detection of cytokines (e.g., *TGFB1* and *TNF*), surface receptors (e.g., *TGFBR* and *CCR4*) and transcription factors (e.g., *STAT6*, and *IRF4*) (P< 0.05, Chi-Square Test, **Figure 1H** and **S2**).

Finally, we performed an additional comparison of enriched human CD4^+^ T cells profiled using Seq-Well S^3 and 10X v2, as well as by Smart-Seq2, a commonly implemented microtiter plate-based approach (**Figure 1F-G**) (Picelli et al., 2013). Integrated analysis of aggregate gene detection revealed that Seq-Well S^3 detects more genes per cell than 10x v2 and nearly as many genes per cell as Smart-Seq2 in pairwise comparison of techniques (10x v2: 2,057 ± 18.7 genes/cell, Seq-Well S^3: 3,514 ± 36.2 genes/cell, SS2: 3,975 ± 74.0 genes/cell; mean ± SEM; P < 0.05, Mann-Whitney Test; **Figure 1F**). Further, comparing the frequency of gene detection between methods revealed crucial differences for transcription factors, cytokines and receptors/ligands. Surprisingly, we observe similar rates of gene detection between S^3 and Smart-Seq2 for a large number of biologically informative genes (**Figure S2F**). Critically, while comparable numbers of genes were detected across methods, Seq-Well S^3 detected more genes per aligned read than either 10x v2 or SS2 in pairwise comparisons (P<0.05, Mann-Whitney U Test; **Figure 1G**).

### A Resource of Cellular States Across Healthy and Inflamed Skin

To demonstrate the utility of Seq-Well S^3 to comprehensively describe cellular states across human pathology at unprecedented resolution, we applied it to profile human skin samples spanning multiple, complex inflammatory skin conditions (**Figure 2**) – including acne, alopecia areata, granuloma annulare, leprosy, psoriasis – as well as normal skin (**Figure 2A-B** and **S3A-C**). In total, we processed nine skin biopsies by S^3 and, after data quality filtering, retained 20,308 high-quality single-cell transcriptomes (**Figure 2A-B**).

To examine similarities and differences among these cells across the high-dimensional gene expression space, we selected variable genes, performed UMAP dimensionality reduction, and identified 33 clusters through Louvain clustering in Scanpy (Wolf et al., 2018) (**Figure 2** and **S3A-C**). To collapse clusters to cell-types, we performed enrichment analyses to identify cluster-defining genes (**Figure S3B**) and then manually assigned cell-type identities based on the expression of known lineage markers (**Figure 2C**). We also generated aggregate gene expression profiles and performed hierarchical clustering using a combined list of the top 50 cluster-defining genes for each cluster to further support our annotations and groupings (**Figure S3C**). Ultimately, we recovered a total of 16 primary cell-types, within which there was considerable heterogeneity. The identified cell types include: B cells (marked by expression of *MS4A1* and *CD79A*), dendritic cells (*FCER1G* and *CLEC10A*), endothelial cells (*SELE* and CD93), fibroblasts (*DCN* and *COL6A2*), hair follicles (*SOX9*), keratinocytes (*KRT5* and *KRT1*), macrophages (*CD68* and *CTSS*), mast cells (*CPA3* and *IL1RL1*), muscle (*NEAT1* and *KCNQ1OT1*), plasma cells (*IGHG1*), Schwann cells (*SCN7A*), and T cells (*CD3D* and *TRBC2*) (**Figure 2** and **S3A-D**). We next sought to define nuanced cell states within these immune, stromal and parenchymal populations – including T cells, myeloid cells, endothelial cells, dermal fibroblasts, and keratinocytes – across the spectrum of skin inflammation.

### Seq-Well S^3 describes T cell states across inflammatory skin conditions

To determine the range of biological diversity that can be captured using Seq-Well S^3, we first focused on further characterizing T cells across the inflammatory skin conditions examined since each is known to significantly skew T cell phenotypes (**Figure 3**) (Diani et al., 2015; Lowes et al., 2014). We performed dimensionality reduction and sub-clustering across T cells alone (**Figure 3A-B**). Our analysis revealed nine sub-clusters that closely correspond to NK cells and CD8^+^ T cells, as well as several known CD4^+^ T-helper cell (Th) subsets. As before, we used the enhanced sensitivity of S^3 for lineage defining transcripts to help annotate the identity of each sub-cluster; for example, in T cell sub-clusters 5 and 6, respectively, we detected distinct expression of canonical regulatory T cell and Th-17 T cell transcription factors (e.g., *FOXP3* and *RORC*, respectively) and immune receptors (e.g. *TIGIT* and *CXCR6* respectively) (**Figure 3C-E** and **S4**). Additionally, we cross-referenced each sub-cluster’s marker genes against a series of curated signatures in the Savant database (Lopez et al., 2017) to confirm our assignments. This analysis highlighted similarity to previously characterized T cell and NK cell populations (**Figure S4C**).

**Figure 3.**
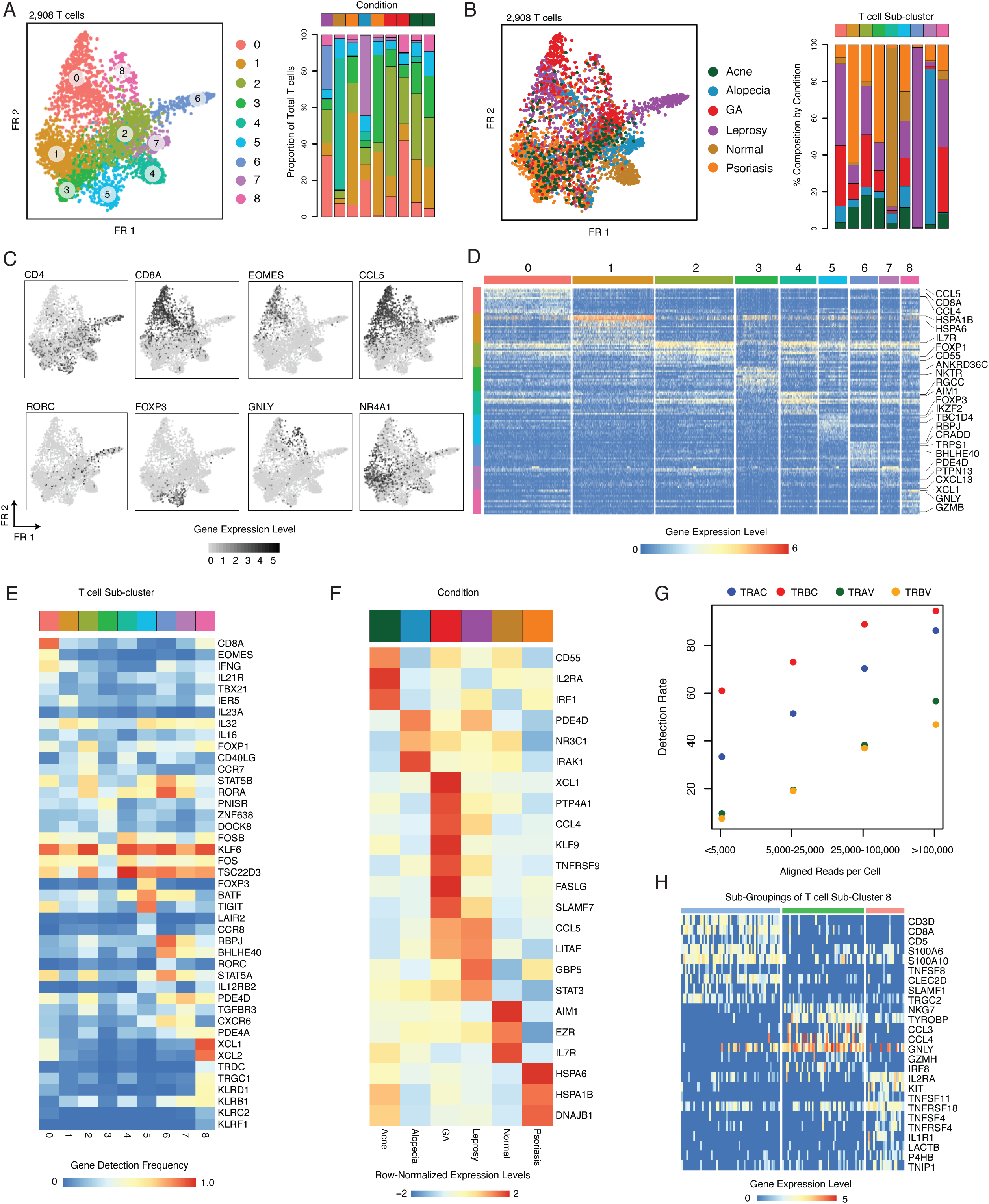
Identification of Inflammatory T cell States using Seq-Well S^3. **A.** (Left) Force-directed graph (Fruchterman Reingold) of 2,908 T cells colored by the nine phenotypic sub-clusters identified by Louvain clustering. (Right) Stacked barplots showing the distribution of these T cell sub-clusters within each skin biopsy. **B.** (Left) Force-directed graph of 2,908 T cells colored by inflammatory skin condition. (Right) Stacked barplots showing the contribution of each inflammatory skin condition to the T cell sub-clusters. **C.** T cell force-directed graphs displaying normalized expression (log(scaled UMI + 1)) of a curated group of sub-cluster-defining gene. Higher expression values are shown in black. **D.** Heatmap showing normalized gene expression values (log(scaled UMI + 1)) for a curated list of sub-cluster-defining genes across nine T cell sub-clusters. **E.** Heatmap showing the rate of detection for lineage-defining transcription factors, cytokines, and cytokine receptors across T cell phenotypic clusters. **F.** Heatmap showing average expression of genes enriched across T cells by inflammatory skin condition (row-normalized average expression values). **G.** Plot showing rates of detection of TCR genes from human skin T cells across a range of sequencing depths. **H.** Heatmap showing normalized gene expression values (log(scaled UMI + 1)) for genes enriched in sub-group analysis of T cell sub-cluster 8.

We next examined T cell phenotypes across inflammatory skin conditions to explore variability in T cell subset composition by skin pathology (**Figure 3B**). This analysis revealed potentially varied contributions to different classes of cutaneous inflammation. For example, sub-cluster 6 is enriched for expression of canonical Th-17 genes including *RORC*, which encodes the Th-17 lineage-defining transcription factor RORγt (Ivanov et al., 2006) and is observed predominantly within the leprosy sample. While either Th1 or Th2 responses are typically thought to predominate in the immune response to leprosy, a role for Th-17 cells in controlling disease has been previously demonstrated (Saini et al., 2013, 2016). We further found that sub-cluster 1, which express *NR4A1*, a transcription factor that is a marker of dysfunctional T cells (Liu et al., 2019), and sub-cluster 3, enriched for genes involved in nuclear organization (*ANKRD36*, *XIST*, and *NEAT1*), were over-represented in both patients from psoriasis (**Figure 3B-C**). In alopecia areata, we detected a unique population of T cells (sub-cluster 7) that overexpress *PDE4D*, which has been shown to plays a role in TCR-dependent T cell activation (**Figure 3D**) (Peter et al., 2007).

We also uncovered considerable variation across cytotoxic T cells and NK cells. Directed analysis within CD8^+^ T cells (sub-cluster 0) revealed a sub-grouping of activated CD8^+^ T cells that express elevated levels of several inflammatory cytokines (*TNF*, *CCL4*, and *XCL1)*, as well as specific affinity receptors (*FASLG* and *TNFRSF9*) and transcription factors (*KLF9* and *EGR2*); this phenotypic skewing was observed primarily in a patient with granuloma annulare (**Figure 3F** and **S4B**). Meanwhile, we found the highest degree of cytotoxic gene expression (*GNLY*, *GZMB*, and *PRF1*) among cells in sub-cluster 8, suggesting that this sub-cluster may represent a diverse set of NK cells, *γδ* T cells, and activated cytotoxic T cells. Indeed, further analysis of sub-cluster 8 revealed 3 distinct component sub-groups of cytotoxic cells: a sub-group of CD8^+^ T cells (T.8.1; *TNFSF8*, *SLAMF1*, *CLEC2D*, *CD5*) expressing various TCR genes; a second sub-group of CD16^+^ cells (T.8.2) expressing cytotoxic effector molecules (*GNLY*, *PRF1*, *GZMB*) and NK surface receptors, consistent with either NK cell or tri-cytotoxic CTL (Balin et al., 2018); and a third sub-group of NK cells (T.8.3) enriched for expression of *c-KIT*, *RANKL* (TNFSF11) and *GITR* (TNFSFR18) (**Figure 3H** and **S4B**) (Söderström et al., 2010).

Profiling of T cell receptor expression is critical to understand T cell antigen specificity (Zhang et al., 2018). Importantly, among CD4^+^ T cells obtained from peripheral blood, we recovered most TCR-V and TCR-J genes at a higher frequency using Seq-Well S^3 as compared to 10x v2 (P< 0.05, Chi-square Test; **Figure S4C**). Among CD4^+^ T cells from peripheral blood, we observed paired detection of TRAC and TRBC in 1,293 of 1,485 CD4^+^ T cells (87.1% Paired Detection Rate, **Figure S4C**). In the setting of skin inflammation, we explored TCR detection rates across a range of sequencing read depths. Overall, we detected *TRAC* in 54.5%, *TRBC* in 75.5%, and paired detection in 46.4% of T cells (**Figure 3G**). Among T cells with at least 25,000 aligned reads, we recovered paired *α* and *β* chains in 66.7%. Among cells from sub-cluster 8, we observe expression of *γ* and *δ* constant genes (*TRGC* and *TRDC*), while remaining T cell clusters exclusively express *α* and *β* TCR constant genes (**Figure S4C**). These data further suggest that sub-cluster 8 represents a diverse population of *γδ*, NK, and cytotoxic CD8^+^ T cells that share common gene expression features and, potentially, roles in inflammation.

### Spectrum of Myeloid Cell States in Skin Inflammation

In the setting of cutaneous inflammation, myeloid cells play a key role in maintaining tissue homeostasis, wound healing and response to pathogens (Malissen et al., 2014). Using Seq-Well S^3, we were able to identify numerous myeloid cell subpopulations defined by a combinations of surface markers, cytokines and lineage-defining transcription factors. Specifically, we independently analyzed 2,371 myeloid cells and identified nine sub-clusters representing 4 primary myeloid cell types based on expression of canonical lineage markers and comparison to cell-type signatures in the Savant database: dendritic cells (*CLEC10A*), Langerhans cells (*CD207* and *CD1A*), macrophages (*CD68* and *CD163*), and mast cells (*CPA3* and *TPSAB1*) (**Figure 4A** and **S4D-E**) (Lopez et al., 2017).

**Figure 4.**
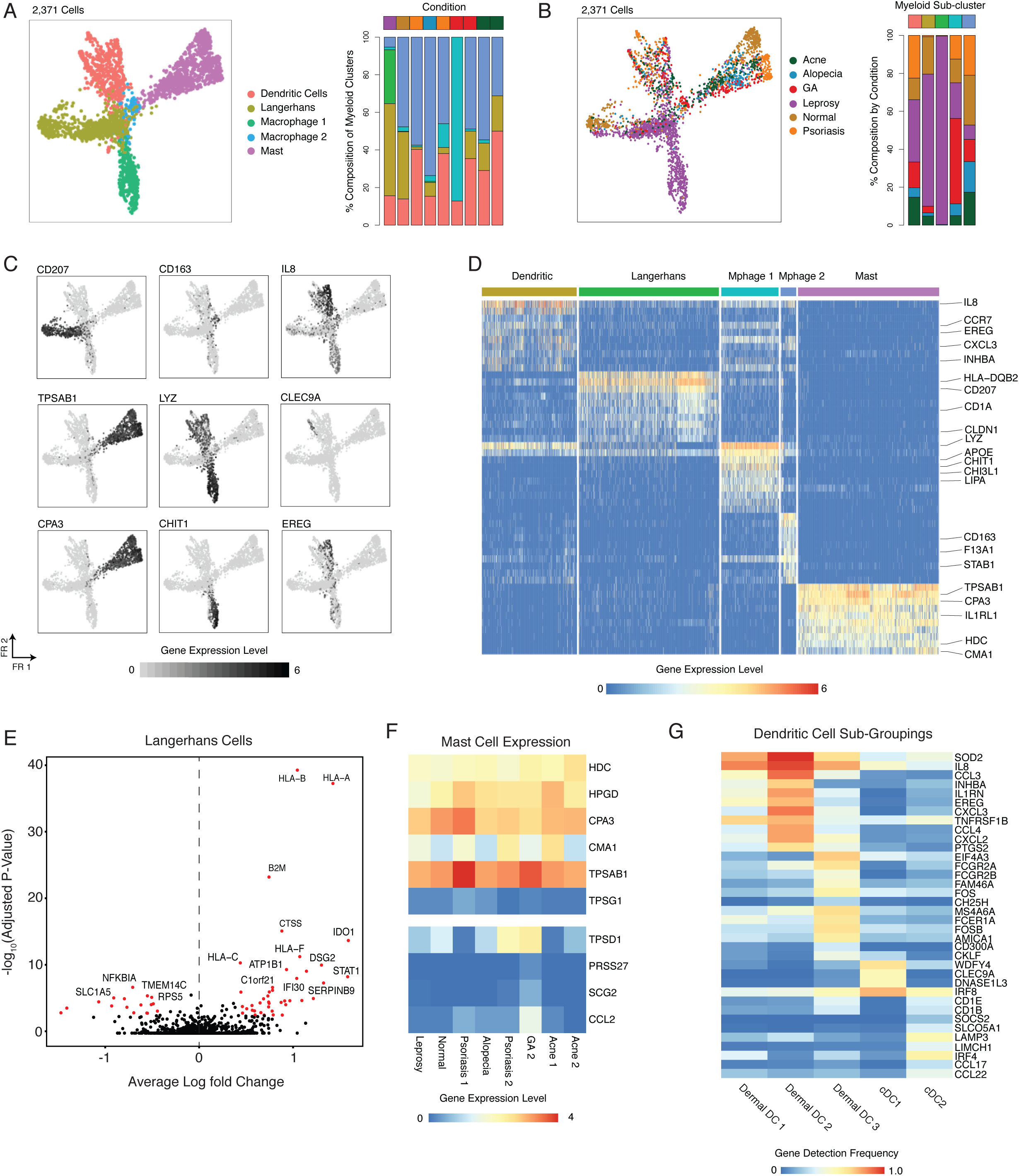
Diverse Myeloid Cell States Uncovered using Seq-Well S^3. **A.** (Left) Force-directed graph of 2,371 myeloid cells colored by five phenotypic sub-clusters (NB, Langerhans cells were enriched from leprosy and normal skin). (Right) Stacked barplots showing the distribution of myeloid sub-clusters within each skin biopsy. **B.** (Left) Force-directed graph of 2,371 myeloid cells colored by inflammatory skin condition. (Right) Stacked barplots showing the contribution of each inflammatory skin condition to each myeloid sub-cluster. **C.** Force-directed graphs of 2,371 myeloid cells that highlighting expression of a curated group of sub-cluster defining genes (log(scaled UMI + 1)). **D.** Heatmap showing the normalized expression (log(scaled UMI + 1)) of a curated list of myeloid cell-type cluster-defining genes. **E.** Volcano plot showing genes differentially expressed in Langerhans cells between leprosy (n_cells_ = 56) and normal skin (n_cells_ = 120). Log10-fold change values are shown on the x-axis and −log10 adjusted p-values are shown on the y-axis. **F.** Heatmaps showing the normalized expression (log(scaled UMI + 1)) of mast-cell proteases across inflammatory skin conditions. **G.** Heatmap showing detection frequencies for transcription factors, surface receptors, and cytokines across DC sub-populations.

Among the macrophages, our data reveal two distinct sub-clusters (**Figure 4A-B**). One macrophage sub-cluster spans normal skin as well as multiple types of skin inflammation and is characterized by elevated expression of previously characterized markers of dermal macrophages (*CD163*, *STAB1*, and *CEPP*) (Fuentes-Duculan et al., 2010). The second sub-cluster, meanwhile, is observed uniquely in leprosy and defined by genes involved in extracellular proteolysis (*LYZ*, *CHIT1*, and *CHI3L1*) (Di Rosa et al., 2013).

Skin functions as both a physical and immunologic barrier, and is the primary site of exposure to environmental antigens. As such, multiple types of antigen-presenting cells (APCs) are distributed in both the dermis and epidermis. In the epidermis, there is a specialized population of antigen-presenting cells known as Langerhans cells. We initially identified Langerhans cells on the basis of expression of canonical markers (*CD207*, *CD1A*; **Figure 4C-D**) (Romani et al., 2003). For biopsies obtained from normal skin and leprosy, we performed MACS enrichments from the epidermal section and loaded Langerhans cells as 5% of the total amount to increase recovery. When we directly compared Langerhans cells from leprosy and normal skin, we observed elevated expression of *IDO1*, *STAT1*, *HCAR3* and MHC class I molecules (*HLA-A*, *HLA-B* and *HLA-F*) in Langerhans cells in leprosy infection, which may suggest a role for Langerhans cells in priming CD8^+^ T cell responses in this disease (**Figure 4E**) (Hunger et al., 2004; Pinheiro et al., 2018).

Additionally, we found a large sub-group of dermal dendritic cells (**Figure 4A**). Further analysis of the *CD207*^−^ dendritic cell sub-cluster revealed multiple sub-groupings of dermal dendritic cells across skin biopsies. Consistent with previous observations from peripheral blood (Villani et al., 2017), we saw a sub-group of dendritic cells that corresponds to cDC1 (*CLEC9A*, *IRF8*, and *WDFY4*) (P<0.05, Permutation Test, **Figure S4H**). We further report another sub-group that represents cDC2 cells (*IRF4*, *SOCS2*, *SLCO5A1, CD1B, CD1E*) (**Figure 4B-C** and **Figure S4F-H**) (Guilliams et al., 2016). Importantly, we detect expression of IL12B, a subunit of the IL-23 cytokine, within the sub-group of IRF4^+^ cDC2 cells (**Figure S4I-J**), which have previously been shown to promote mucosal type 17 inflammation via secretion of IL-23 (Schlitzer et al., 2013). Further, this sub-grouping of cDC2 cells express high levels of *CCL17* and *CCL22*, chemokines involved in T cell chemotaxis (**Figure S4J**) (Stutte et al., 2010).

We further identified three sub-groups of dermal dendritic cells that are broadly distinguished from conventional dendritic cell clusters by expression of *CLEC10A* (**Figure S4J**), which has been shown to influence T cell cytokine responses in skin (Kashem et al., 2015; Kumamoto et al., 2013). Cells from dermal DC sub-group 1 show elevated expression of *CD44*, *IL8* and *SOD2* (**Figure S4I**). Cells from dermal DC sub-group 2 display elevated expression of pro-inflammatory chemokines up-regulated during DC maturation (*CXCL3*, *CCL2* and *CCL4*) (Jin et al., 2010) and soluble mediators (*EREG* and *INHBA*). Finally, a third sub-grouping of dermal DCs (Dermal DC3) was distinguished by expression of *FCER1A*, *FCGR2A*, and *FCGR2B*, which are important for interfacing with humoral immunity (**Figure S4I**) (Guilliams et al., 2014).

In the skin, mast cells are most commonly associated with allergic responses, but mast cell proteases serve additional roles in inflammation and pathogen defense (Pejler et al., 2010). Among skin mast cells, we detect core expression of *HDC* (Histidine decarboxylase), *HPGD*, and *TPSAB1* (Tryptase *α*/*β* 1) (**Figure 4F**) (Dwyer et al., 2016). Importantly, we observe variable expression of mast cell proteases *TPSD1* (Tryptase D1) and *CMA1* (Chymase A1), which are primary mast cells effector molecules (Pejler et al., 2010), which may have functional consequences. By performing analysis across inflammatory conditions and patients, we identify a distinct pattern of mast cells with elevated expression of proteases (*TPSD1*, Tryptase D1 and *PRSS27*, serine protease 27), *SCG2* (secretogranin 2), and *CCL2* in a patient with granuloma annulare (**Figure 4F**).

### Detection of Endothelial Heterogeneity and Vascular Addressin Expression

Multiple types of endothelial cells exist within the dermis of the skin. As in most tissues, arterioles shuttle oxygenated blood to tissues terminating in a capillary bed that gives rise to post-capillary venules. Importantly, DARC^+^ post-capillary venules are the primary site of egress of immune cells from circulation into tissues (Schön et al., 2003). Using the improved sensitivity of Seq-Well S^3, we sought to understand the spectrum of endothelial cell diversity and vascular addressin expression across multiple instances of skin inflammation (von Andrian and Mempel, 2003). We performed sub-clustering and dimensionality reduction across 4,996 endothelial cells (**Figure S5A-B**) and identified three primary sub-clusters of dermal endothelial cells defined by distinct expression patterns: vascular smooth muscle (*TAGLN*), endothelial cells (*CD93*) and lymphatic endothelial cells (*LYVE1*) (**Figure S5C**). Importantly, we found multiple sub-clusters of CD93^+^ endothelial across normal and inflamed skin biopsies (**Figure S5A-B**). For example, we observe two distinct populations of endothelial cells: a population of DARC^−^, CD93^+^ endothelial cells (Venule sub-cluster 3) that displays elevated expression of *SLC9A3R2*, which is involved in endothelial homeostasis (Bhattacharya et al., 2012), and another cluster of proliferating endothelial cells (Venule sub-cluster 4) (**Figure S5D**).

Further, we sought to understand the distribution of vascular addressins expressed by DARC^+^ endothelial sub-populations, the site primary site of lymphocyte egress into tissues (**Figure S5E**) (Thiriot et al., 2017). Notably, across sub-populations of CD93^+^ endothelial cells (Venule sub-clusters 1-4), we observe variation in expression of vascular addressins (**Figure S5E**). Among post-capillary venules, we observe broadly elevated expression of *ITGA5*, *ITGA6*, *ITGB4*, *ICAM2*, and *ITGA2*, while arterioles express higher levels of *ITGA7*, *ITGA8*, and *ITGB5*. Further, we observe the highest expression of *ITGA4*, *ITGA9*, *ITGB2* and *ITGB8* among lymphatic endothelial cells (**Figure 5E**).

**Figure 5.**
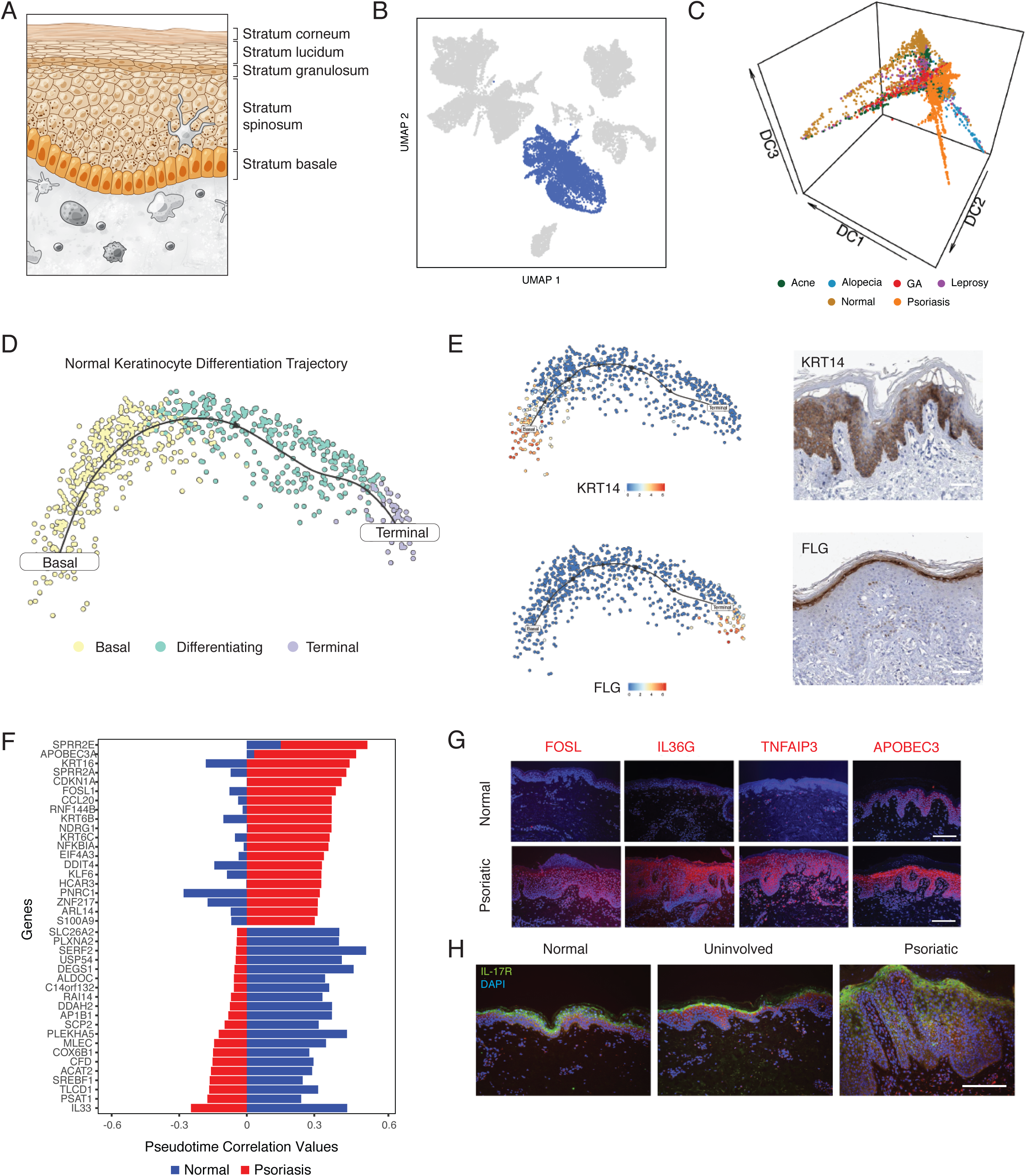
Keratinocyte Differentiation Trajectories. **A.** Diagram showing the layers of the epidermis and morphologic changes associated with keratinocyte differentiation. **B.** UMAP embedding of 20,308 cells with all keratinocyte and hair follicle populations highlighted in blue. **C.** Diffusion map of 5,141 keratinocytes colored by inflammatory skin condition. Axes correspond to diffusion components. **D.** t-SNE plot showing differentiation trajectory of keratinocytes from normal skin from basal cells (yellow) through differentiating cells (aqua) and terminal keratinocytes (purple). **E.** (Top-left) tSNE plot of normal keratinocytes with normalized *KRT14* expression values overlayed. (Top-right) Immunohistochemistry staining showing the expression of *KRT14* from the human protein atlas (Uhlén et al., 2015). (Bottom-left) tSNE plot of normal keratinocytes with normalized *FLG* expression values overlayed. (Bottom-right) Immunohistochemistry staining of *FLG* from the human protein atlas(Uhlén et al., 2015). Scale bars = 50 microns. **F.** Stacked barplot showing genes with the highest differential pseudo-time correlation between normal keratinocytes (blue) and psoriatic keratinocytes (red) sorted by correlation values in psoriatic keratinocytes. Correlation values shown on the x-axis represent Pearson correlation coefficients between normalized gene expression and diffusion pseudotime. **G.** (Top) Immunofluorescence staining in normal (above) and psoriatic (below) for FOSL, IL36G, TNFAIP3, and APOBEC3. All images stained for nuclei (DAPI) and gene of interest (Red Fluorescence). Scale bar = 100 microns. **H.** Immunofluorescence staining for IL-17R expression (green) in normal (left), uninvolved (middle), and psoriatic skin (right). Scale bar = 100 microns.

### Altered Dermal Fibroblast Identities in Skin Inflammation

Dermal fibroblasts provide structural support and are the primary source of extracellular matrix components within the skin. Previous studies have demonstrated significant variation among dermal fibroblasts based on their relationship to anatomic features of the skin (Driskell and Watt, 2015; Driskell et al., 2013). To deeply catalogue diverse fibroblast cell states across inflamed skin, we performed dimensionality reduction and sub-clustering within the 4,189 fibroblasts identified across all samples and conditions (**Figure S5F-G**). In comparison to inflamed biopsies, fibroblasts from normal skin display enrichments in *LTBP4*, *IGFBP5*, and *TCF4*. Consistent with previous single-cell studies of dermal fibroblasts, we observe a sub-population of fibroblasts (Cluster 6) that express *COL11A1*, *DPEP1* and *RBP4*, where these cells were suggested to have a role in connective tissue differentiation (**Figure S5H**) (Tabib et al., 2018).

Fibroblasts from GA patient 1 (sub-cluster 2) express elevated levels of *SPOCK1*, *CRLF1*, and *COMP*, a cartilage protein that is upregulated in matrix-producing fibroblasts following myocardial infarction (Fu et al., 2018) (**Figure S5H-I**). Further, fibroblasts from GA patient 2 (sub-cluster 0) display elevated expression of protease inhibitor 16 (*PI16*), which inhibits the function of MMP2 (Hazell et al., 2016), and *ITIH5*, a protease inhibitor important for maintenance of dermal hyaluronic acid that is overexpressed in skin inflammation (**Figure S5H-I**) (Huth et al., 2015). Finally, among fibroblasts from acne patients, we observed elevated expression of multiple metallothioneins (**Figure S5H-I**). Specifically, the expression levels of *MT1E* and *MT2A* are highest in fibroblasts and endothelial cells in acne (**Figure S5H**). As seen among endothelial cells, fibroblast expression patterns in acne are consistent with a wound healing response (Iwata et al., 1999).

### Keratinocyte Differentiation Trajectories

Within the epidermis, keratinocytes undergo a stereotyped differentiation process in which cells acquire altered morphology and phenotype as they mature (**Figure 5A**) (Fuchs, 1990). Under physiologic conditions, basal keratinocytes are characterized by expression of *KRT14* and *TP63*, and continuously divide to give rise to the remaining cells of the epidermis (Fuchs and Raghavan, 2002). Using keratinocytes from normal skin, we performed pseudo-temporal analysis to reconstruct the differentiation process of normal epidermal keratinocytes (**Figure 5D**). More specifically, in normal skin, we first identified a population of keratinocytes enriched for expression of *TP63* and *KRT14*, markers of basal keratinocytes (**Figure S6B**) (Pellegrini et al., 2001). We then used known patterns of cytokeratin expression to infer localization of keratinocytes along a supervised differentiation trajectory (**Figure 5E** and **S6A**) (Ordovas-Montanes et al., 2018). Our trajectory analysis revealed patterns of transcription factor and cytokeratin expression that closely correspond to previously established signatures of keratinocyte maturation (Cheng et al., 2018). Consistent with immunohistochemical staining from the Human Protein Atlas (**Figure 5E**) (Uhlén et al., 2015), we observed enrichment of filaggrin (*FLG*), a protein in the outer layers of the epidermis (Sandilands et al., 2009), *mRNA* among keratinocytes that lie at the terminal points in the pseudo-temporal ordering (**Figure 5E** and **Figure S6B**).

Having established a trajectory for normal keratinocyte differentiation, we next examined patterns of keratinocyte differentiation across pathologic conditions. To identify conserved and unique patterns across conditions, we constructed a combined diffusion map using the 5,141 keratinocytes recovered across all samples (**Figure 5C**). While keratinocytes from most conditions closely align with normal differentiation, we observe marked deviation in the differentiation trajectory of psoriatic keratinocytes (**Figure 5C**). Consistent with previous observations, differential expression analysis reveals significant up-regulation of antimicrobial peptides (*S100A7*, *S100A8*, *S100A9*) and pro-inflammatory cytokines (*IL36G*, *IL36RN*) in psoriatic keratinocytes (Li et al., 2014).

Based on increased sensitivity of Seq-Well S^3 to detect transcription factors observed in peripheral lymphocytes, we hypothesized that our data might enable identification of novel transcriptional regulators of psoriatic keratinocytes. To identify potential drivers of the psoriatic disease process within the epidermis, we performed differential pseudo-time correlation analysis between psoriatic and normal keratinocytes. Specifically, we separately constructed pseudo-time trajectories for normal and psoriatic keratinocytes, calculated correlation values between diffusion pseudo-time and gene expression levels, and examined the difference in correlation values between psoriatic and normal keratinocytes (**Figure 5F** and **S6A-B**). Notably, we observed positive correlation of *FOSL1*, an AP-1 transcription factor, with diffusion pseudo-time in psoriatic keratinocytes, implying that *FOSL1* is preferentially expressed along the differentiation trajectory of psoriatic keratinocytes. To validate this observation, we performed immunofluorescence staining for FOSL1 protein, and measured increased levels of FOSL1 in psoriatic skin (**Figure 5G**). We further validated the distribution of additional genes overexpressed or differentially correlated with diffusion pseudo-time in psoriatic keratinocytes (including *TNFAIP3, IL36G*, and *APOBEC3*) at the protein level (**Figure 5G** and **S6A**).

To further define differences in gene expression patterns between normal and psoriatic keratinocytes, we scored the expression levels of known cytokine response signatures using a series of reference signatures gene lists derived from population RNA-Seq of cultured keratinocytes exposed to IL-17A (**Figure S6C**). While IL-17 has been previously implicated in the pathogenesis of psoriasis, here we infer the identity of cells that dominate the IL-17 response, localizing the expression of IL-17 responsive genes to spinous keratinocytes (Ordovas-Montanes et al., 2018). To validate this observation, we performed immunofluorescent staining for IL-17R protein and measured the highest staining within spinous keratinocytes exclusively within psoriatic skin (**Figure 5H**). Collectively, these data provide novel insights into the localization IL-17 response in psoriatic keratinocytes.

## DISCUSSION

Here, we present an enhanced technique for high-throughput scRNA-Seq – Seq-Well S^3 – that affords improved sensitivity for transcript capture and gene detection. Through use of a templated second-strand synthesis, S^3 recovers information typically lost in bead-based high-throughput scRNA-Seq protocol such as Seq-Well or Drop-Seq. Specifically, S^3 reclaims mRNA molecules that are successfully captured and reverse transcribed but not labeled with a second primer sequence through template switching (**Figure 1** and **S1**). Using Seq-Well S^3, we obtain a 5-10 fold increase in the number of unique molecules captured from cells at similar sequencing depth relative Seq-Well v1 (**Figures 1**, **S1** and **S2**) (Gierahn et al., 2017). Beyond aggregate increases in the number of transcripts recovered per-cell, the improvements in sensitivity made possible by Seq-Well S^3 enable enhanced detection, and thus deeper examination, of lineage-defining factors in immune and parenchymal cells – such as transcription factors, cytokines, and cytokine receptors among lymphocytes (**Figure 1** and **S2**) – which are often transiently or lowly expressed (Zhu et al., 2010). Among CD4^+^ T cells isolated from PBMCs, for example, we observed rates of gene detection similar to those observed in Smart-Seq2, a best-in-class microtiter plate-based method (**Figure 1F-G** and **S2F**).

Similarly, using Seq-Well S^3, we report improved paired detection of *α* and *β* TCR sequences from T cells in peripheral blood and tissue biopsies (**Figures 3G** and **S4C**). Among CD4^+^ T cells from PBMCs, we recover paired TCR *α* and *β* constant genes in 87.1% of cells. Together with targeted enrichment, amplification and sequencing, we anticipate that Seq-Well S^3 will enable improvements in TCR reconstruction and deep characterizations of clonotype-phenotype relationships at scale (Zhang et al., 2018). Collectively, our validation experiments show that Seq-Well S^3 significantly augments the amount of information that can be recovered in massively-parallel scRNA-seq experiments, enabling high-resolution profiling of low-input biopsy samples at scale.

With this enhanced method, here, we move towards a draft atlas of human skin inflammation by creating a compendium of cell-types and states for the broader research community (Regev et al., 2018). Through use of Seq-Well S^3, we survey, at unprecedented resolution, the diversity of cell-types and states – e.g., among tissue resident T cells and myeloid cells – present across multiple types of skin inflammation. For example, GA and leprosy are two granulomatous diseases characterized by aggregates of lymphocytes and macrophages within the dermis, which are both thought to arise from a delayed-type hypersensitivity response to *M. leprae* infection (leprosy) and an unknown agent (GA) (Modlin et al., 1984; Terziroli Beretta-Piccoli et al., 2018). Here, we find that both are characterized by the presence of T cell sub-cluster 0 (Immature CD8^+^ CTL) and T cell sub-cluster 8 (mature CTL effectors containing CD8^+^ T-CTL, *γδ* and NK cells; **Figure 3**). Although both conditions contain CD163^+^ dermal macrophages and various DC subpopulations, M1-like macrophages were present only in leprosy, which host the intracellular pathogen *M. leprae*, were present only in leprosy (Fulco et al., 2014; Verreck et al., 2004). Moreover, GA uniquely contained specific populations of fibroblasts expressing *SPOCK1*, *CRLF1*, and *COMP* (**Figure S5**), which likely reflect remodeling of the dermis with mucin deposition and alternation of elastin fibers (Piette and Rosenbach, 2016; Yun et al., 2009).

Acne, meanwhile, is an inflammatory disease thought to arise in response to infection with *P. acnes*, resulting in the formation of lesions that resemble a wound following eruption of the hair follicle into the dermis (Beylot et al., 2014). Here, we observe 2 clusters of endothelial cells marked by expression of *SLC9A3R2*, a marker of endothelial homeostasis, and a signature of proliferation (**Venule clusters 3 and 4**, **Figure S5**). This increased angiogenesis and endothelial proliferation is most consistent with the proliferative phase of wound healing in acne (Holland et al., 2004).

Alopecia areata and psoriasis both arise from autoimmune and autoinflammatory processes, yet there were distinct differences in their underlying cell states. For example, alopecia areata is thought to be driven by a population of CD8^+^ T cells that target hair follicles (Xing et al., 2014). Notably, in alopecia, we report a sub-cluster of T cells characterized by expression of *PDE4D* (**Figure 3**). PDE4 inhibitors have recently shown demonstrated efficacy in the treatment of alopecia (Keren et al., 2015; López et al., 2017), and it is intriguing to speculate that these inhibitors might work by targeting this subset of T cells.

In psoriasis, T cells are thought to be a primary driver of inflammation, with dendritic cells playing a central role in the recruitment and polarization of T cells that contribute to the hyperproliferation of keratinocytes in the disease (Lowes et al., 2014). In both patients with psoriasis, we report a sub-cluster of DCs (IRF4^+^ cDC2) that display elevated expression of *CCL17*, *CCL22* and *IL12B* (**Figure 4G** and **Figure S4I**). Importantly, a similar population of dermal cDC2 cells has recently been shown to drive psoriatic inflammation in mice and humans through the recruitment of inflammatory T cells (Kim et al., 2018; Zaba et al., 2010). Although we detected a diversity in T cell subtypes in psoriatic lesions, we note few Th-17-like cells (Hawkes et al., 2018).

Leveraging the increased sensitivity of Seq-Well S^3, we performed pseudo-time correlation analysis to uncover an altered differentiation trajectory of keratinocytes compared to normal skin (**Figure 5** and **S6**). From our pseudo-time correlation analysis, we detected FOSL1 as a putative transcription factor involved in psoriatic differentiation, a finding which we validated through immunofluorescent staining of healthy and psoriatic skin (**Figure 5G**). Further, previous studies using *in vitro* keratinocyte based systems have suggested that more differentiated keratinocytes were the main responders to IL-17A, given larger effect sizes in differentiated compared to monolayer keratinocyte (Chiricozzi et al., 2014). Using data generated with Seq-Well S^3 cross-analyzed against an IL-17 response signature in keratinocytes, we show that IL-17 responses are observed in keratinocytes from all layers of the epidermis, but that these responses are stronger in keratinocytes derived from more differentiated layers of the psoriatic epidermis (**Figure S6C**). This observation is corroborated by co-localization of the IL-17 receptor subunits (IL-17RA/IL-17RC) in the upper layers of psoriatic epidermis (**Figure 5H**).

In conclusion, we describe a powerful massively-parallel scRNA-Seq protocol that enables improved transcript capture and gene detection from low-input clinical samples. Here, Seq-Well S^3 provides novel insights into putative mechanisms and the cellular localization of previously appreciated and unknown responses to specific inflammatory mediators in immunologic skin conditions in a fashion not previously achievable. Increases in the sensitivity of gene and transcript detection are increasingly important as single-cell atlasing efforts shift from detection of large differences between cell types within normal tissue to identification of subtle differences in cell state across cell types within diseased tissues. The increased sensitivity of gene detection and transcript capture afforded by S^3 enhances the strength of inferences that can be drawn from these types of single-cell data, as evidenced by the range of immune, stromal and parenchymal cell states uncovered in human skin inflammation. The S^3 protocol is easy to integrate into current bead-based RNA-Seq platforms, such as Drop-Seq, making it broadly useful for the single-cell community, particularly in the setting of human disease. Importantly, S^3’s increases in library complexity and sequencing efficiency reduce costs relative to plate-based protocols, and providing researchers with a powerful and cost-effective alternative to commercial solutions in a format that can be deployed almost anywhere.

## ACKNOWLEDGEMENTS

This work was supported in part by the Koch Institute Support (core) NIH Grant P30-CA14051 from the National Cancer Institute, as well as the Bridge Project, a partnership between the Koch Institute for Integrative Cancer Research at MIT and the Dana-Farber/Harvard Cancer Center. This work was also supported by the Food Allergy Science Initiative at the Broad Institute and the NIH (5P01AI039671, 5U19AI089992). A.K.S. was supported by the Searle Scholars Program, the Beckman Young Investigator Program, the Pew-Stewart Scholars Program for Cancer Research, a Sloan Fellowship in Chemistry, the NIH (1DP2GM119419, 2U19AI089992, 2R01HL095791, 1U54CA217377, 2P01AI039671, 5U24AI118672, 2RM1HG006193, 1R33CA202820, 1R01AI138546, 1R01HL126554, 1R01DA046277, 1U2CCA23319501) and the Bill and Melinda Gates Foundation (OPP1139972, OPP1202327, OPP1137006, and OPP1202327). R.L.M. was supported by the NIH (R01 AI022553, AR040312, AR074302). J.G. is supported by the Taubman Medical Research Institute and the NIH (R01-AR060802, R01-AI30025, and P30-AR075043). T.K.H. is supported by the NIH F30-AI143160.

## CONTRIBUTIONS

**T.K.H., M.W.H., T.M.G., R.L.M., J.C.L., and A.K.S.** designed the study. **T.K.H., M.H.W., T.D., D.W., P.A.**, and **B.A.** collected skin samples and performed single-cell sequencing experiments. **S.S., L.C.T., and J.E.G.** performed immunofluorescent staining. **T.K.H., M.H.W., T.M.G., and F.M.** analyzed data under the guidance of **J.O.M., R.L.M., J.C.L., and A.K.S**. **T.K.H., M.H.W., T.M.G., J.C.L., and A.K.S.** wrote the manuscript with input from all authors.

## DECLARATION OF INTERESTS

A.K.S, and J.C.L. have received compensation for consulting and SAB membership from Honeycomb Biotechnologies. A.K.S. has received compensation for consulting and SAB membership from Cellarity, Cogen Therapeutics, and Dahlia Biosciences. T.M.G., T.K.H., M.H.W., A.K.S., and J.C.L are co-inventors on a provisional patent application filed by MIT relating to the improved methodology described in this manuscript.

---

**Supplementary Figure 1.**
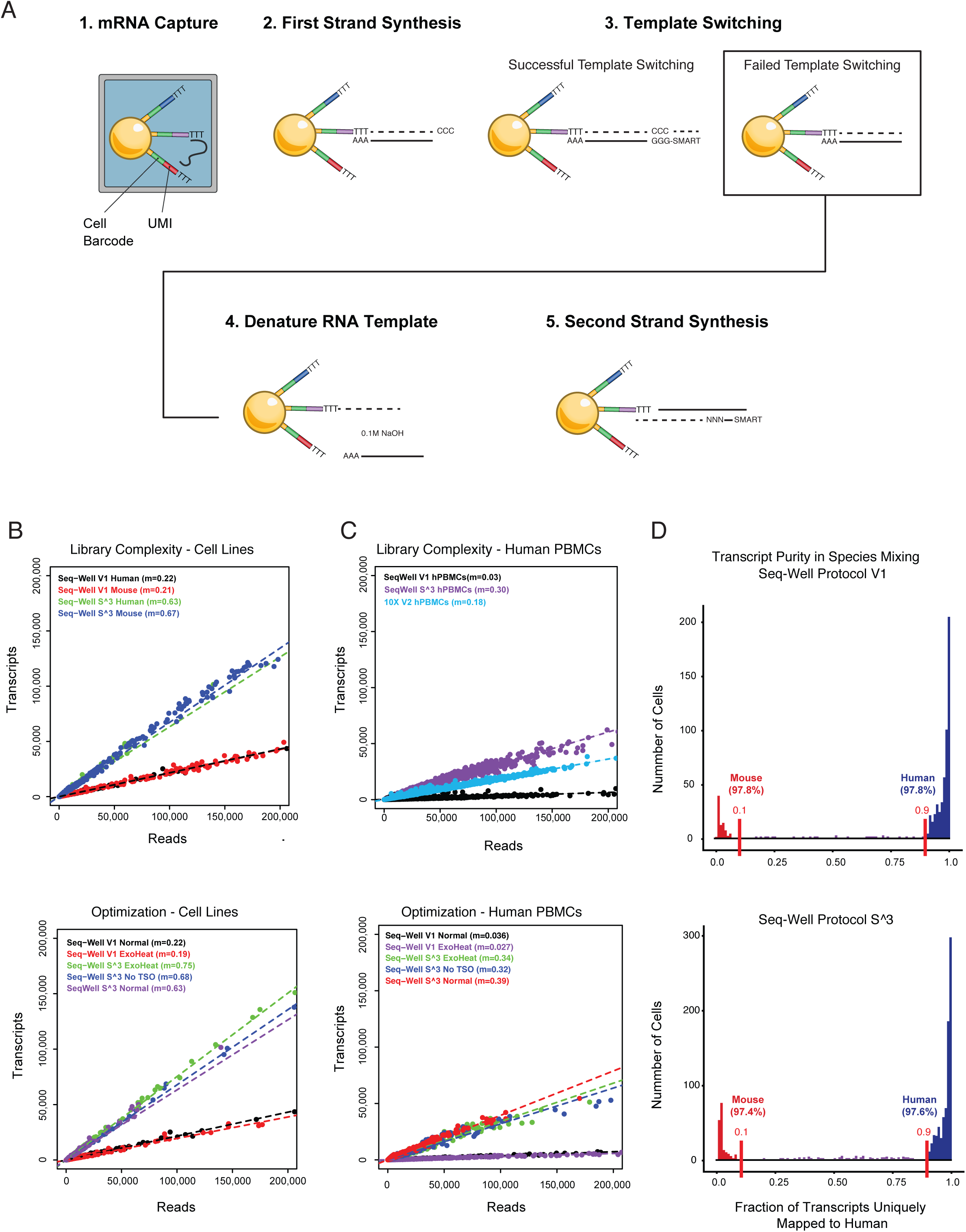
Second-Strand Synthesis Overview, related to Figure 1. **A.** Illustration of the second strand synthesis procedure: (1) mRNA is captured via poly-T priming of poly-adenylated mRNA; (2) First strand synthesis is performed to generate single-stranded cDNA template on bead-bound sequences; (3) Successful template switching: The use of enzymes with terminal transferase activity generates a 3’ overhang of 3 cytosines. Template switching utilizes this overhang to append the SMART sequence to both ends of the cDNA molecule during first strand synthesis. Failed Template Switching: If template switching fails, this results in loss of previously primed and reverse transcribed mRNA molecules; (4) mRNA template is chemically denatured using 0.1M NaOH; (5) Second strand synthesis is performed using a random-octamer with the SMART sequence in the 5’ orientation; and, (6) Following second strand synthesis, PCR amplification, library preparation and sequencing are performed to generate data. **B.** Scatterplots show the relationship between transcript detection (y-axis) and number of aligned reads per cell (x-axis) for an initial experiment (top) series of optimization conditions using HEK293 and NIH-3T3 cell lines (botttom). **C.** Scatterplots that illustrate the relationship between number of transcripts detected (y-axis) and number of aligned reads per cell (x-axis) between Seq-Well V1 and Seq-Well S^3 in sequencing experiments for an initial experiment (top) and a series of optimization experiment using human PBMCs (bottom). **D.** Histograms that show the fraction of transcripts uniquely mapped to the human genome for each cell for Seq-Well V1 (Top) and Seq-Well S^3 (Bottom). Colors indicate species classification for cells with at least 90% purity of human (blue) or mouse (red) mapping.

**Supplementary Figure 2.**
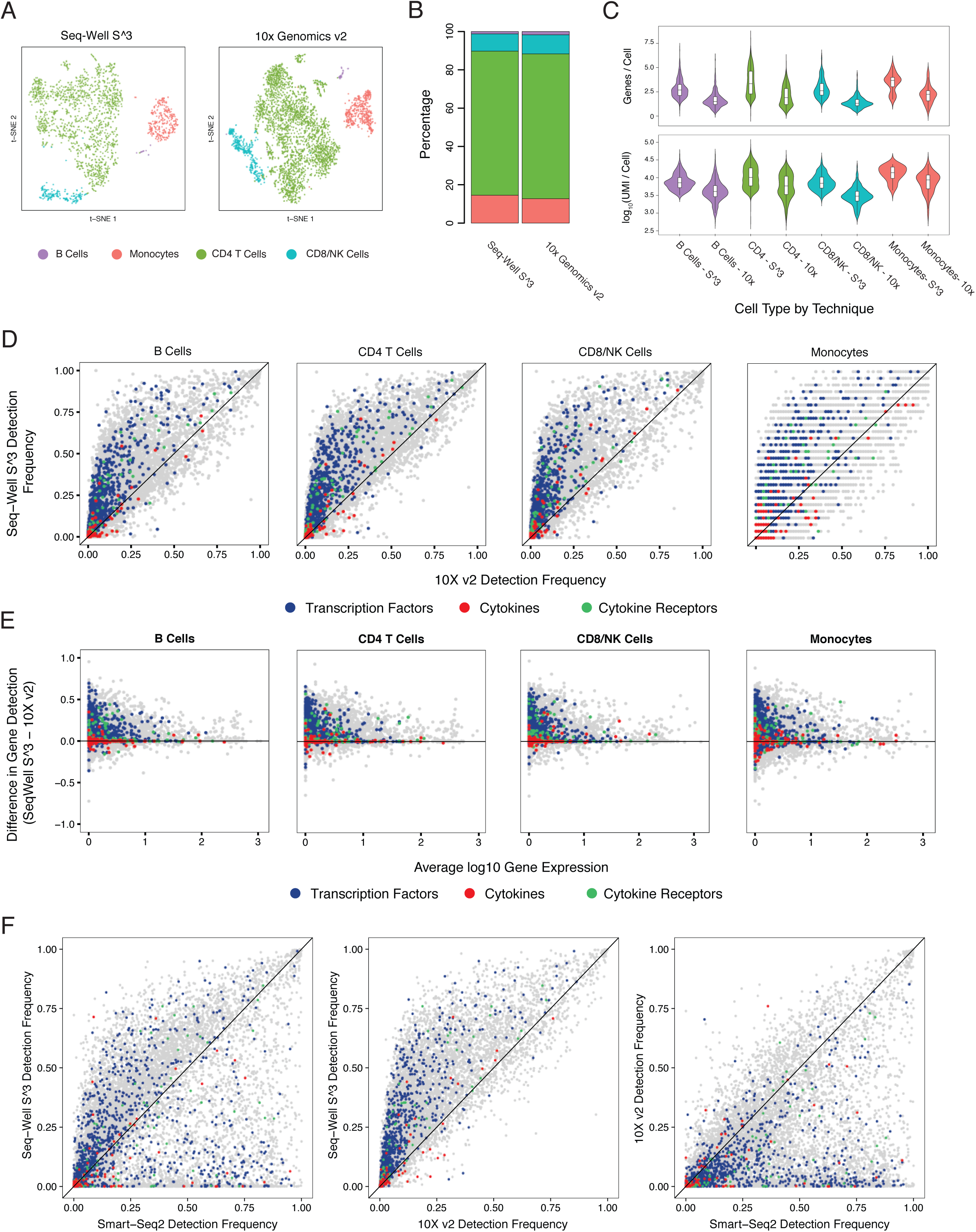
PBMC Methods Comparisons, related to Figure 1. **A.** t-SNE plot showing detected cell-types among PBMCs including CD4^+^ T cells (green), CD8^+^/NK Cells (blue), B cells (purple), and Monocytes (red) using 10X v2 and Seq-Well S^3. Cells recovered using Seq-well are colored with darker shades. **B.** Stacked barplots show the proportion of cell types recovered using Seq-Well S^3 (left) and 10X v2 (right). **C.** Top: Violin plots (boxplots median +-quartiles) showing the distribution of per cell gene detection from Seq-Well S^3 (left) and 10X v2 (right). Bottom: Violin plots (boxplots median +/− quartiles) showing the distribution of per cell-gene detection from Seq-Well S^3 (left) and 10X v2 (right). **D.** Scatterplots showing a comparison of gene detection frequencies between Seq-Well S^3 (y-axis) and 10x v2 (x-axis) for each cell type. **E.** Scatterplots showing the difference in gene detection between Seq-Well S^3 and 10X v2 (y-axis) as a function of average normalized expression (log(scaled UMI + 1)) (x-axis). **F.** Scatterplots showing a comparison of gene detection frequencies among sorted CD4^+^ T cells between **(Left)** Seq-well S^3 (y-axis) and 10x v2 (x-axis), (**Middle)** Seq-Well S^3 (y-axis) and Smart-Seq2 (x-axis), and **(Right)** 10x v2 (y-axis) and Smart-Seq2 (x-axis).

**Supplementary Figure 3.**
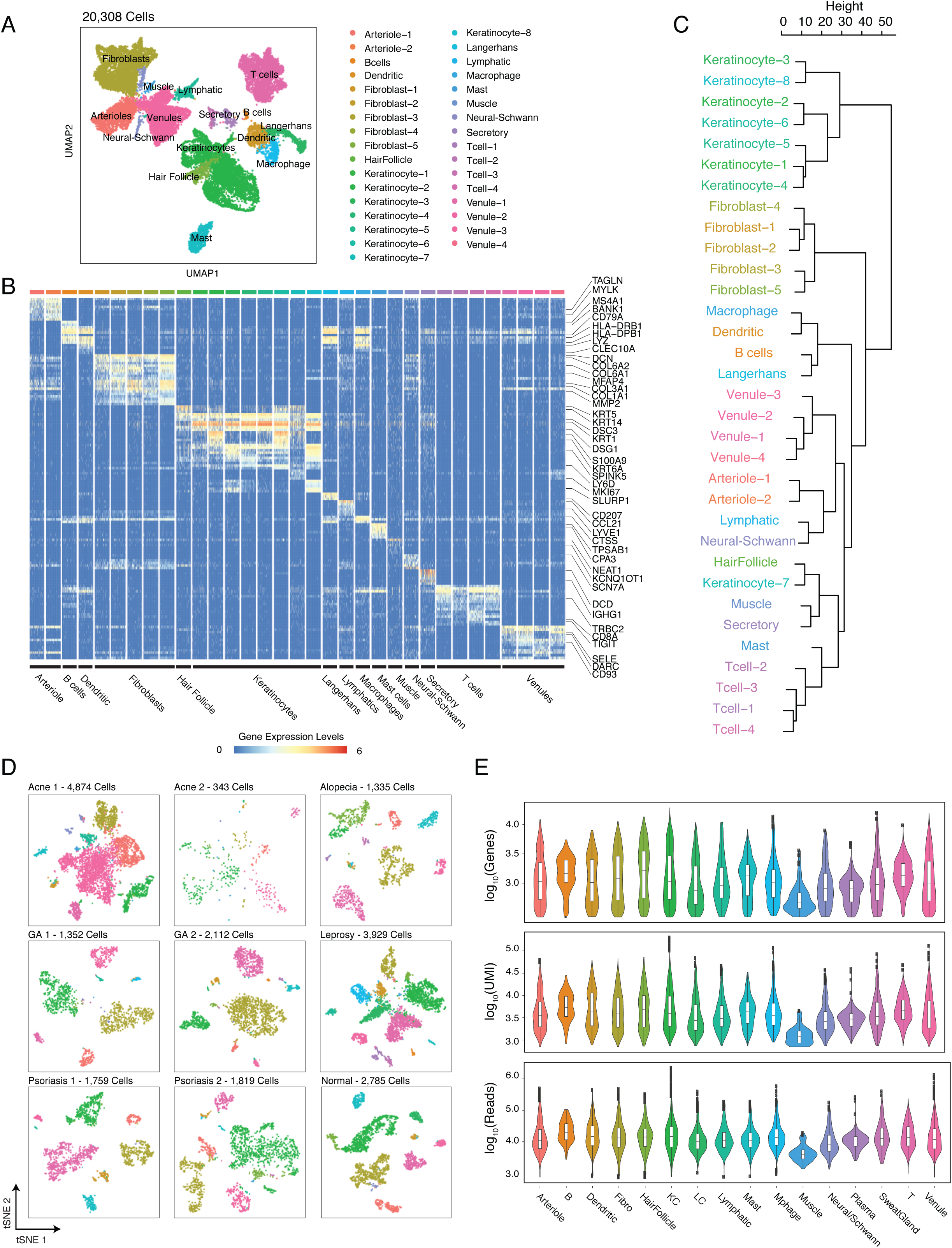
Overview of Samples, related to Figure 2. **A.** UMAP plot for 20,308 cells colored by 33 cell type cell type clusters (Louvain Resolution: 2.0). **B.** Heatmap showing the relative expression of cell-type defining gene signatures across 20,308 cells. **C.** Dendrogram of hierarchical clustering shows similarity of cell type clusters among top 25 cluster-defining genes (**Figure S3B**). **D.** t-SNE plots for each of the nine skin biopsies colored by generic cell type. **E.** Violin plots show the distribution of per-cell quality metrics displayed in UMAP embedding of 20,308 cells colored by colored generic cell-type classification (Figure 2B).

**Supplementary Figure 4.**
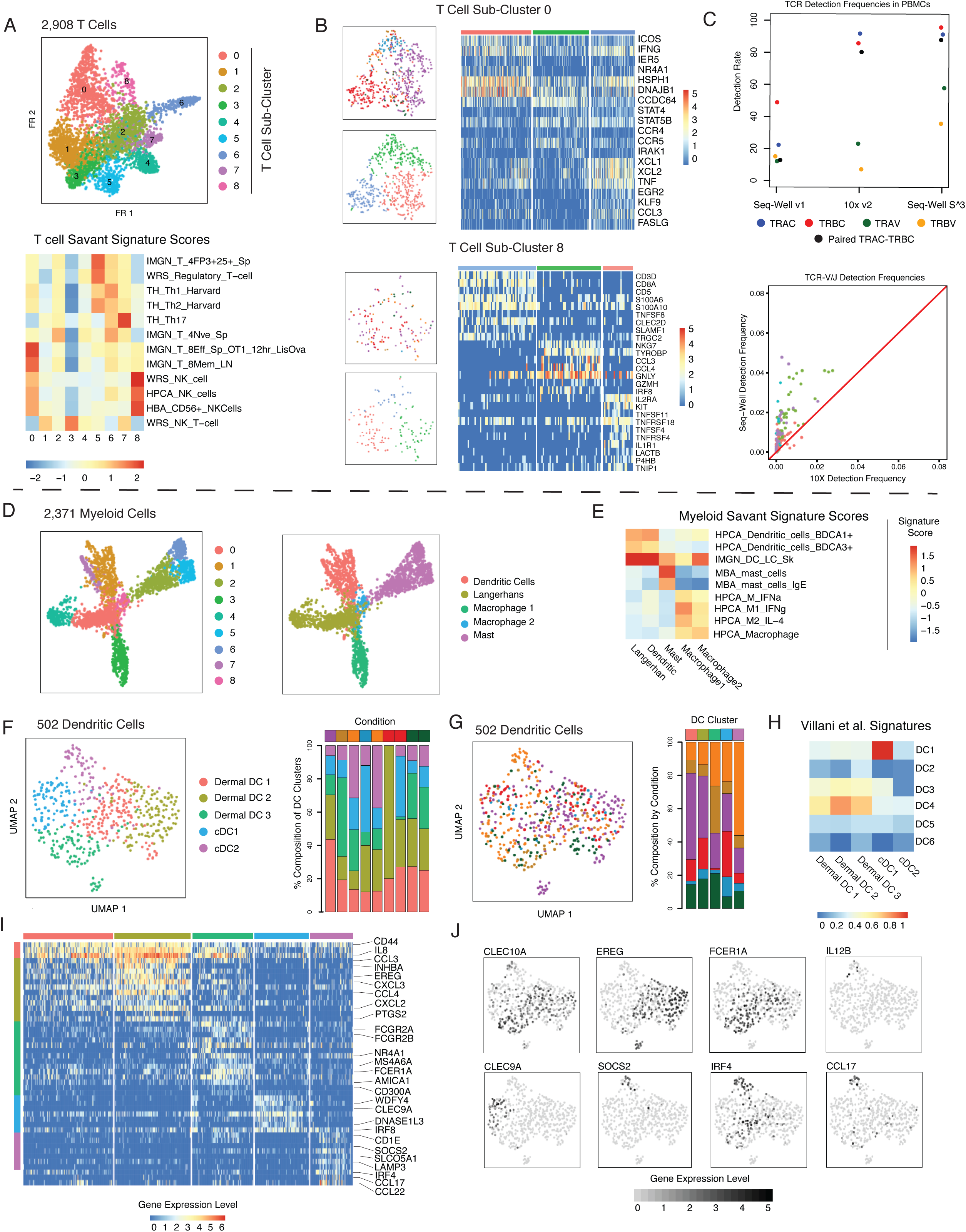
Immune Cell Heterogeneity, related to Figures 3 and 4. **A.** (Top) Force-directed graph of 2,903 T cells colored by T cell sub-cluster. (Bottom) Heatmap of gene-set enrichment scores based on comparison of T cell phenotypic sub-clusters to a curated list of reference signatures in the Savant database. **B.** Sub-grouping results for (top) T cell sub-cluster 0 and (bottom) T cell sub-cluster 8. For each analysis, t-SNE plots colored by inflammatory skin condition (top-left) and sub-cluster (bottom-left) are shown. For each clusters, heatmaps show gene expression patterns across T and NK cells sub-types (right). **C.** (Top) Detection rates for TCR genes for PBMCs in Seq-Well v1, 10x v2. and Seq-Well S^3. (Bottom) Detection frequency of TCR V-J (e.g. TRAV/J and TRBV/J) genes in CD4^+^ T cells from peripheral blood between Seq-Well S^3 (y-axis) and 10x v2 (x-axis). Colors correspond to TRAJ (red), TRAV (green), TRBJ (blue), and TRBV (purple) genes. **D.** Force-directed graph of 2,371 myeloid cells colored by myeloid phenotypic sub-clusters. **E.** Heatmap of gene-set enrichment scores based on comparison of myeloid phenotypic sub-clusters to a curated list of reference signatures in the Savant database. **F.** (Left) UMAP plot for 502 dendritic cells from human skin colored by phenotypic sub-grouping. (Right) Stacked barplot showing composition of dendritic cells within each of nine skin biopsies by DC sub-cluster. **G.** (Left) UMAP plot for 502 dendritic cells from human skin colored by inflammatory skin condition. (Right) Stacked barplot showing contribution of inflammatory skin conditions to each dendritic cell sub-grouping. **H.** Heatmap showing average signature score across 5 dermal DC populations based on dendritic cell signatures from *Villani et al. Science 2017*. **I.** Heatmap showing the distribution of normalized gene expression levels (log(scaled UMI + 1)) for cluster-defining genes across dermal DC subpopulations. **J.** UMAP plots colored by normalized expression levels for DC sub-grouping-defining genes.

**Figure S5.**
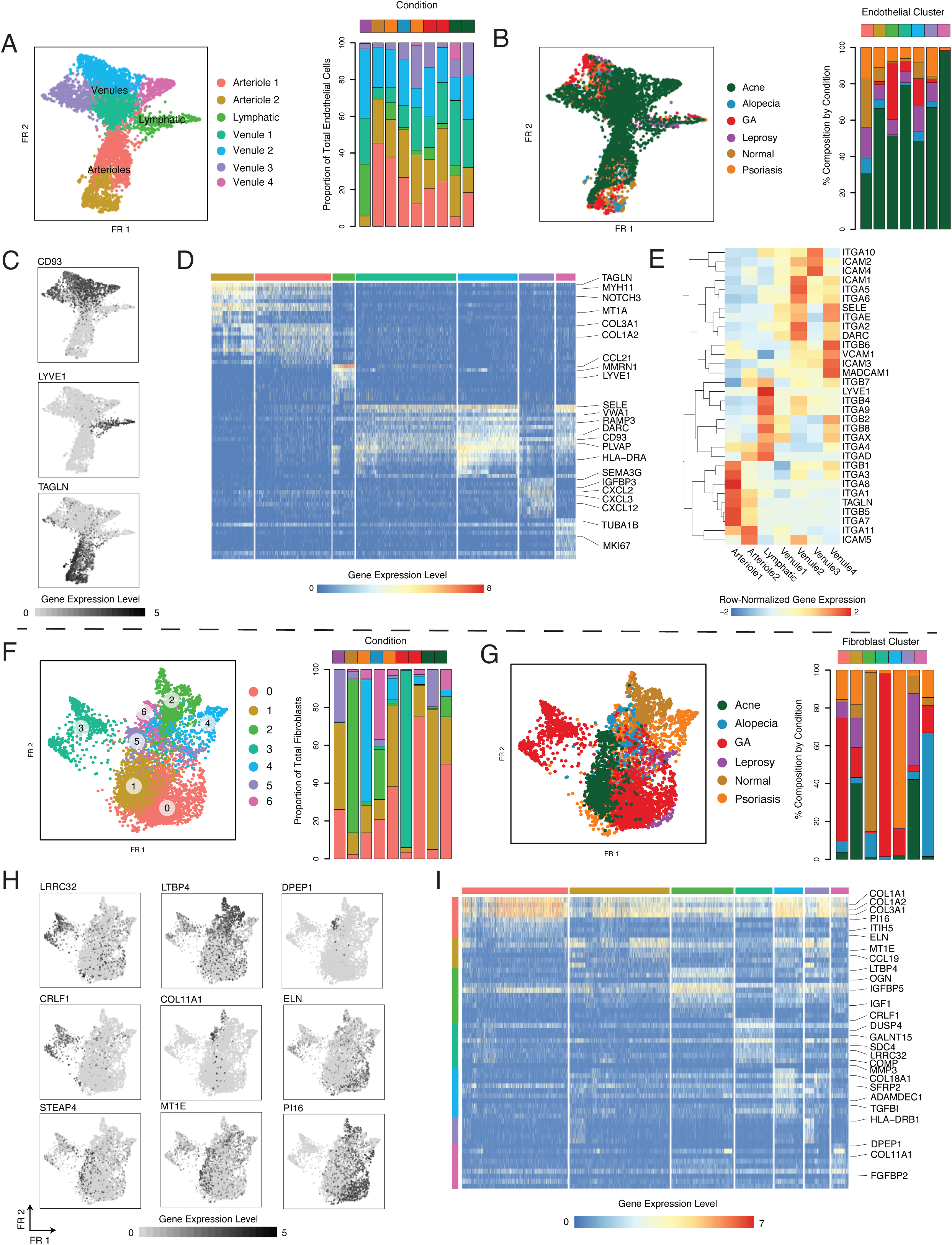
Stromal Cell Diversity. **A.** Force-directed plots for 4,996 endothelial cells colored by phenotypic sub-cluster (left) and stacked barplot showing the distribution of endothelial phenotypic sub-clusters across samples (right). **B.** Force-directed plots for 4,996 endothelial colored by inflammatory skin condition (left) and stacked barplot showing the contribution of each inflammatory skin condition to endothelial phenotypic sub-clusters. **C.** Forced-directed plot colored by normalized expression level of genes that mark endothelial cell types: (Left) CD93, venules, (Middle) TAGLN, arterioles, (Right) LYVE1, lymphatics. **D.** Heatmap showing patterns of normalized gene expression levels (log(scaled UMI + 1)) across 7 clusters of endothelial cells. **E.** Heatmap showing row-normalized expression levels of vascular addressins across phenotypic sub-clusters of endothelial cells. **F.** Force-directed plots for 4,189 fibroblasts colored by phenotypic sub-cluster (left) and stacked barplot showing the distribution of fibroblast phenotypic sub-clusters across samples (right). **G.** Force-directed plots for 4,189 fibroblasts colored by inflammatory skin condition (left) and stacked barplot showing the contribution of each inflammatory skin condition to fibroblast phenotypic sub-clusters. **H.** Force-directed graphs highlighting fibroblast cluster defining genes. **I.** Heatmap showing the normalized gene expression levels (log(scaled UMI + 1)) of fibroblast cluster-defining genes.

**Supplementary Figure 6.**
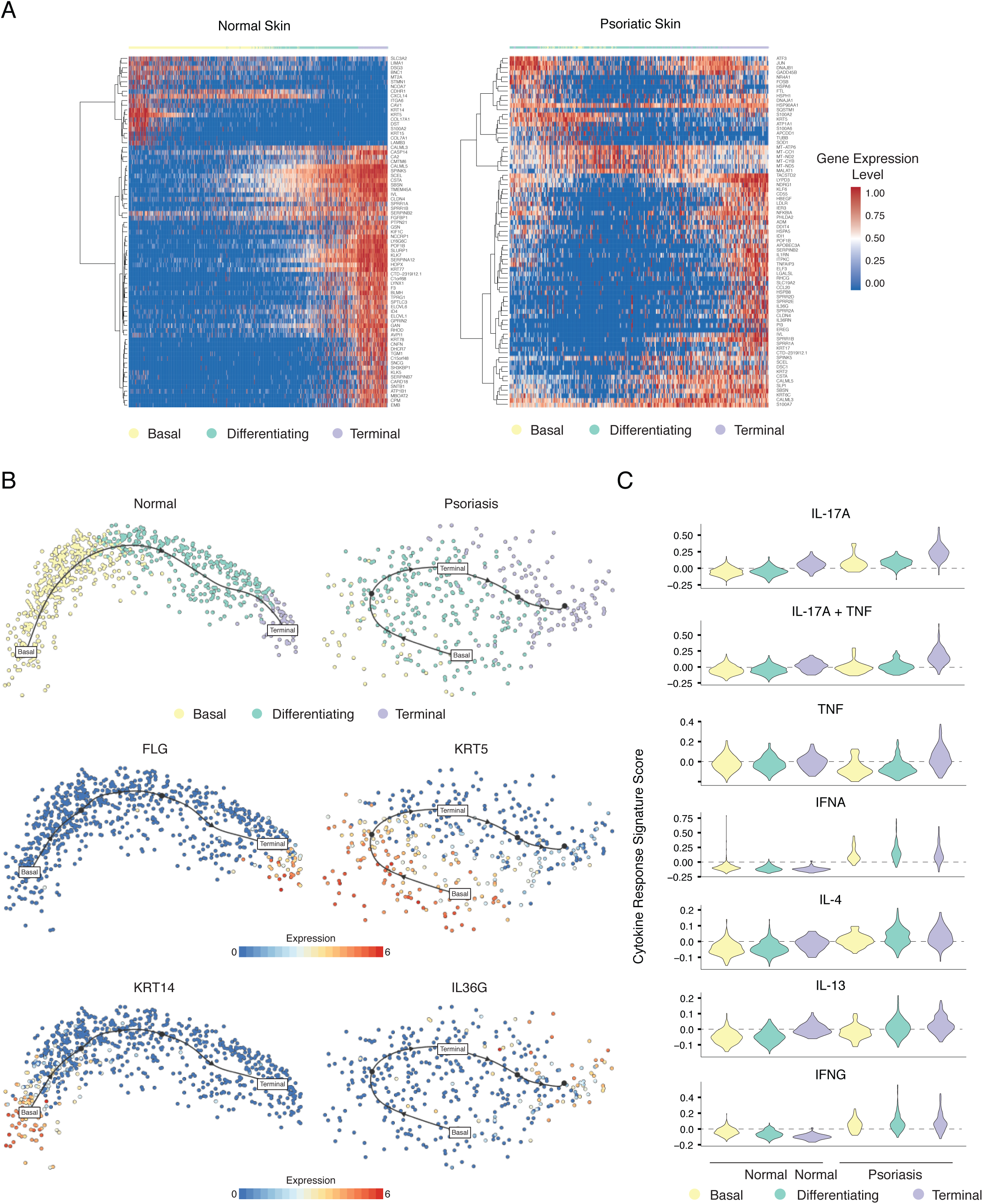
Keratinocyte Differentiation Trajectories, related to Figure 5. **A.** (Left) Heatmap showing enrichment of genes along pseudo-temporal trajectories for normal keratinocytes. (Right) Heatmap showing enrichment of genes along pseudo-temporal trajectories among psoriatic keratinocytes. **B.** Differentiation trajectories for Normal (left) and Psoriatic (right) keratinocytes. **C.** Violin plots showing localization of cytokine response signatures in basal, differentiating and terminal keratinocytes for Normal (left) and Psoriatic (right) keratinocytes.

